# Reinforcement Learning for Control of Human Locomotion in Simulation

**DOI:** 10.1101/2023.12.19.572447

**Authors:** Andrii Dashkovets, Brokoslaw Laschowski

## Abstract

Control of robotic leg prostheses and exoskeletons is an open challenge. Computer modeling and simulation can be used to study the dynamics and control of human walking and extract principles that can be programmed into robotic legs to behave similar to biological legs. In this study, we present the development of an efficient two-layer Q-learning algorithm, with k-d trees, that operates over continuous action spaces and a reward model that estimates the degree of muscle activation similarity between the agent and human state-to-action pairs and state-to-action sequences. We used a human musculoskeletal model acting in a high-dimensional, physics-based simulation environment to train and evaluate our algorithm to simulate biomimetic walking. We used imitation learning and artificial biomechanics data to accelerate training via expert demonstrations and used experimental human data to compare and validate our predictive simulations, achieving 79% accuracy. Also, when compared to the previous state-of-the-art that used deep deterministic policy gradient, our algorithm was significantly more efficient with lower computational and memory storage requirements (i.e., requiring 7 times less RAM and 87 times less CPU compute), which can benefit real-time embedded computing. Overall, our new two-layer Q-learning algorithm using sequential data for continuous imitation of human locomotion serves as a first step towards the development of bioinspired controllers for robotic prosthetic legs and exoskeletons. Future work will focus on improving the prediction accuracy compared to experimental data and expanding our simulations to other locomotor activities.

## I. Introduction

Control of robotic leg prostheses and exoskeletons is an open problem and an active area of research [1], [2]. Computer simulation can be used to study the dynamics and control of human walking and extract principles that can be programmed into robotic legs to behave similar to biological legs. These models should ideally be fully-predictive such that the controller captures the underlying mechanisms of locomotion. Studies have used musculoskeletal models to simulate and optimize walking with passive exoskeletons [3], [4] and for knee osteoarthritis rehabilitation [5]. Predictive simulation of human locomotion has mainly been driven by optimization, specifically optimal control theory [6]-[9].

Optimal control involves solving an optimization problem to determine the control signals that maximize performance. For locomotion, optimal control can find control policies that generate walking to meet objectives such as stability or energy efficiency [9]. The controller addresses the optimization problem by minimizing a function (e.g., metabolic energy expenditure or summed muscle activations). However, designing an objective function that represents natural gait dynamics requires significant domain expertise. Studies have also used computed muscle control to estimate muscle activations that generate motion that follows a desired trajectory [10], [11]. However, this method only reproduces a predefined trajectory and cannot predict responses to new inputs.

In addition to optimal control, reinforcement learning can be used to simulate the dynamics and control of human locomotion [12]-[15]. In reinforcement learning, an agent learns to make decisions by interacting with an environment, receiving feedback through rewards, and adjusting its behavior to maximize cumulative reward. An advanced version is deep reinforcement learning, which uses deep neural networks. Deep reinforcement learning has been used to train controllers for autonomous walking robots [16]-[18] and exoskeletons [19]-[22]. Compared to optimal control, reinforcement learning tends to require less human input to manually tune the controllers and are more adaptable to learning novel tasks. However, as concluded in a recent review on deep reinforcement learning for simulation of human locomotion control [23], “better controllers could be trained by utilizing imitation learning for a set of optimal motions”. Imitation learning, a class of reinforcement learning, involves training an agent to mimic or imitate the behavior of an expert by learning from demonstrations rather than relying solely on rewards or explicit instructions.

In this study, we present the development of a new two-layer Q-learning algorithm with k-d trees and sequential data optimization to train a learning agent to imitate human walking based on expert demonstrations. Our algorithm features a new combined reward model, which first estimates the degree of muscle activation similarity between the agent and human state-to-action pairs. In the case of data presence of state-to-action sequences, it additionally amends the score to improve stability. We were able to simulate biomimetic walking with a 79% similarity compared to experimental human data [24]. Moreover, when compared to the previous state-of-the-art that used deep deterministic policy gradient to simulate human walking [23], [25], [26], our algorithm was significantly more efficient with lower computational and memory storage requirements (i.e., requiring 7 times less RAM and 87 times less CPU compute), which can benefit real-time embedding computing for robotic leg control.

## II. Methods

### A. Generation of Artificial Walking Data

We first generated artificial walking data. The goal of this step is to generate human-like data that describes the states and muscles activations that need to be done for each state in order to transfer to the next state to walk. To generate human-like data, we used the OpenSim simulation environment and the OpenSim musculoskeletal controller [27], [28]. We used the software to model the human musculoskeletal system, derive the equations of motion, and synthesize the movements by integrating them over time. The environment builds on another open-source project, Simbody, for physics-based simulation. The OpenSim musculoskeletal controller contains predefined rules that describe human gait dynamics. When launching the application, our program generates new instances of the OpenSim environment, the OpenSim rule-based controller, and k-d tree memory buffer and starts generating artificial data for training. Fig. 1 shows the collection of synthetically generated data of human-like walking from interactions with the OpenSim rule-based controller and environment. Using this approach, we are able to increase in our research capabilities as our solution becomes less dependent on real human testing, thus saving time and money associated with experimental data collection.

**Fig. 1.**
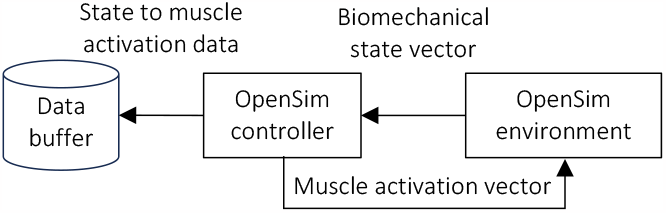
Data generation of biomechanical states and muscle activations using the OpenSim simulation environment.

Our human musculoskeletal model consists of 18 musculotendon actuators, nine in each leg, including biarticular hamstrings, biceps femoris, gluteus maximus, iliopsoas, rectus femoris, vasti, gastrocnemius, soleus, and anterior tibialis [29], [30]. The muscle force is dependent on its length, velocity, and activation, which is the control signal to the muscle that generates force and can vary from 0% to 100%. Our model also includes ligaments and ground contact forces. Ligaments are biological structures that generate force when stretched beyond a predetermined length. For the hip, knee, and ankle joints, the ligaments were modeled as rotational springs with increasing stiffness as the joint angle increased. Foot-ground contact was modeled using a compliant Hunt-Crossley contact model. Two contact spheres were located at the heel and toes, and forces were generated based on the depth and velocity of the spheres penetrating the ground.

The OpenSim environment calculates and returns states every 10 milliseconds over the 10-second simulation. At each iteration, the agent (i.e., the system that interacts with the environment) receives the observed state vector, which includes 1) joint angles and angular velocities of the pelvis, hip, knee, and ankle joints, and 2) positions and velocities of the pelvis, center of mass, head, torso, toes, and talus. The controller outputs a vector of current muscle excitations based on the observation vector or internal states, and current strength. At each iteration, the agent returns a vector of 22 muscle excitations.

Our algorithm initializes the musculoskeletal model based on a predefined biomechanical state vector, describing the initial posture of the human model from which walking begins. The state vector gets delivered to the OpenSim controller, which proceeds the current state vector per defined rule-based logic and outputs the vector of muscle activations. The muscle activation action vector is stored in a memory buffer with the current state vector. The environment accepts the action vector and executes the action as per the action vector values. Once the action is performed, the environment returns the new state vector to the controller. This process repeats for 50 episodes to complete one epoch. During this process, we collect the initial set of state-action pairs to use in environment exploration.

### B. Q-Learning with K-dimensional Trees

The first layer of our algorithm generates single-state bio-mechanical vectors and selects the most optimal muscle activation vectors for that state using the reward function. We use model-free Q-learning to determine an optimal policy for each continuous action. The term model-free refers to algorithms that do not explicitly build a model of the environment but rather learn from interactions with the environment without trying to understand or represent its dynamics. Since we used vector-based states and actions, standard Q-learning would struggle to create appropriate Q-tables, which is a data structure that stores the action-value function, known as the Q-function, for each state-action pair in a Markov decision process. The Q-table is used to guide the agent’s decision-making by providing an estimate of the expected cumulative reward for taking an action in a particular state. The Q-table is often represented as a 2D table, where each row corresponds to a scalar state and each column corresponds to a scalar action. The value in each cell represents the estimated cumulative reward, called the Q-value, for choosing that action in that state. The Q-values are updated as the agent explores the environment and learns from experiences. The agent can then choose the action with the highest Q-value in each state to maximize its long-term cumulative reward. However, the Q-table can become large in high-dimensional environments, such as in our case. It is also difficult to use Q-tables in cases where the states and actions are represented as vectors. Thus, we used the idea of Q-learning as the foundation to our algorithm to store decision-making data in a format that enables easy interpretation of results. This approach usually requires fewer computational resources compared to deep neural networks. Also, function approximation and policy gradient methods may not be ideal in this case as they do not provide a direct means to interpret and adjust results.

We decided to estimate the Q-function using two k-dimensional trees with a rewards dictionary [31]. K-d trees offer quick nearest-neighbour queries for state and action vectors as they provide a balanced and efficient representation of multi-dimensional data, enabling fast search operations in high-dimensional spaces. We used k-d trees, specifically the sklearn KDTree model, for Q-learning with continuous values instead of using Q-table for discrete values.

The agent training process is shown in Fig. 2. The agent is represented as the instance that contains k-d trees for the state vectors, action vectors, and the rewards dictionary, which get used as per previously generated state-action pairs. We obtain new state vectors and generate the action vector based on the rule-based OpenSim controller. We send the current state vector to our agent to generate another action vector. The agent selects the action vector by searching the nearest state vectors inside the existing k-d tree and returns its index. We obtain the previously related action vector to this state and search for similar action vectors. We select the action vector with the highest reward based on values from our rewards dictionary.

**Fig. 2.**
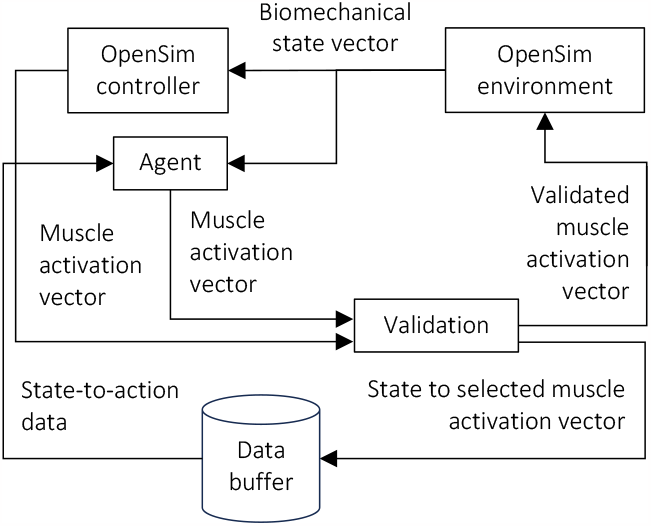
Data generation of biomechanical states and muscle activations via training our reinforcement learning agent with action validation.

Once both action vectors are generated by the agent and OpenSim rule-based controller, the agent vector validation process starts. This validation process aims to ensure that the agent generates the valid action vector for the current state and prohibits its later use in the case of a poor action vector. This helps increase the robustness of our learning agent. To validate the action vector, we calculate the cosine distance between the action vectors from the agent and OpenSim controller. If the vector differs by more than 0.1, we reject the action.

In reinforcement learning, the agent needs manage exploration and exploitation trade-off [32], which involves finding an optimal balance between exploring new actions and exploiting already learned knowledge to maximize cumulative rewards. In other words, we can choose an action that is known to be rewarding (exploitation) or try a new action that might be more rewarding (exploration). Focusing solely on exploitation may lead to suboptimal performance if the agent gets stuck in a locally optimal policy without exploring other options. However, excessive exploration can lead to inefficiencies or delays in discovering the optimal policy if the agent keeps exploring without sufficiently exploiting already learned knowledge [32]. The exploration coefficient is the desired trade-off between exploration and exploitation, and typically ranges between 0 and 1, where 0 represents no exploration (pure exploitation) and 1 represents complete exploration (pure random action selection).

We used imitation learning during exploration, which does not require a high coefficient compared to deep reinforcement learning [32]. Unlike deep reinforcement learning, where exploration is crucial for discovering optimal policies, in imitation learning, the goal is to imitate an expert’s behavior or policy. Although exploration is not the main focus of imitation learning, it can still be beneficial. For example, if the expert demonstrations are limited or imperfect, incorporating some exploration can help the agent handle novel situations not previously covered by the expert. Since our expert (i.e., the Open-Sim rule-based controller) generates action vectors for ideal states and does not provide information how to handle postural deviations, we cannot set this value to 0. Since we have high-dimensional vectors with environment variability, we set the coefficient to 0.1. We chose this coefficient arbitrarily based on the 3-sigma rule. Initially, we tested 2 sigma, 95% confidence interval, and a coefficient value of 0.05. However, the model training was unstable. We decided to reduce the confidence interval to 90% and set the coefficient value to 0.1. After validation, we send the agent’s valid action vector to the environment to obtain the next state vector or send the action vector generated by the OpenSim controller.

### C. Agent Optimization with Sequential Data

Once the first layer is trained, we acquired knowledge on how our agent should behave in particular state. At this stage, the agent can receive the current state vector, search for the nearest state vector in the memory buffer, and find the muscle activation vectors that were previously used for the nearest state vector. However, we do not know which muscle activation vector for the nearest state vector usually leads to the longest sequence of state-to-action pairs. Here, we assume that the longer sequence, the less likely the human model will fall and thus successfully walk. Consequently, we developed a second layer with the goal of determining the optimal muscle activation vector from the first layer based on the length of state-to-action sequences. Thus, the second layer of our algorithm uses sequential data and the same implementation of a tree search to learn a trajectory that defines the sequence of continuous actions (Fig. 3). This method leverages the strengths of the idea of Q-learning, which is effective at learning optimal policies for discrete actions, and tree search over sequential data, which is effective at learning optimal sequences of actions.

**Fig. 3.**
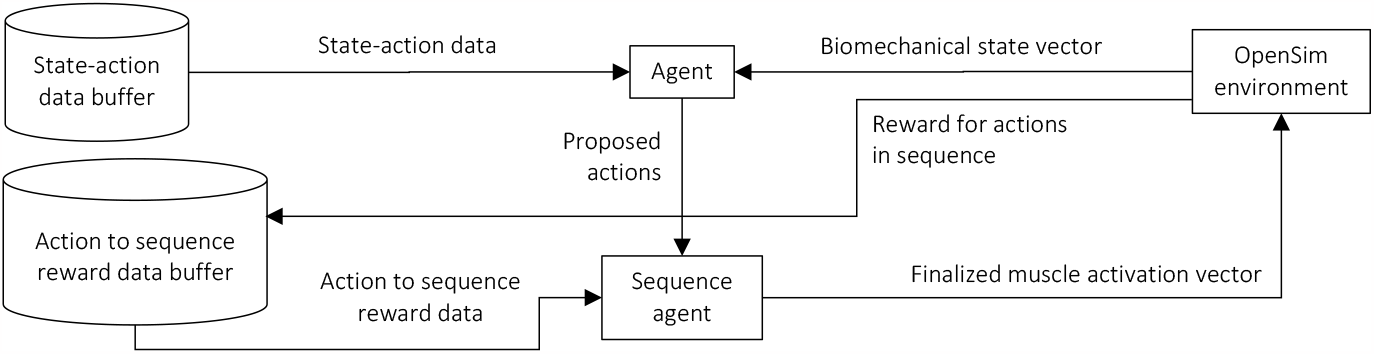
Second layer of our reinforcement learning agent training based on sequential data optimization.

We initially experimented with a simple reinforcement learning model to learn to walk with low computational requirements. However, the simulation would get stuck in the middle of the gait cycle, where the agent tried to maintain the same state as the last posture. This occurred because the environment returned higher rewards in this state compared to rewards in states that lead to more human-like walking. The agent became stuck in the local maximum of our imitated function and was not able to neglect this reward and continue walking. This issue is known as the delayed rewards problem where rewards received by an agent are delayed or sparse, making it challenging for an agent to associate actions with long-term outcomes. To address this problem, we developed a second layer to our algorithm to consider both the reward for a given state and the reward for the entire sequence of state-action pairs over a given epoch.

We trained our agent to learn from sequences of state-action pairs and accumulate experiences based on sequential rewards. Here we use the term “learn” because the second layer does not acquire information during the previous stages. Instead, it receives information from experiments conducted by the first layer and adjusts them over time by gathering them into sequences.

After the first layer of our algorithm, the agent obtains knowledge and can make decisions on how to best behave in each state. Instead of selecting the action via reward maximization, our agent returns the best three actions from the state-action reward and sends them to the sequence agent (i.e., the second layer), which selects the best action by a sequence reward function. Unlike the state-action reward generated by the environment, we custom-designed a reward function based on averaging the lengths of state-action sequences. The sequence agent selects the action from the proposed actions via reward maximization based on the average length of sequences where this action is present. This reward function assumes that the agent should not select an action that presents itself in short sequences as these sequences often lead to deficient performance.

If there is no data in the sequence reward data buffer or the sequence lengths of actions are equal, we do not use sequence optimization, the agent selects the action based on the state-action reward and sends the final muscle activation vector to the OpenSim environment. At the end of the epoch, we calculate the length of state-action pairs in the completed sequence and send this information to the action to sequence reward data buffer. This learning agent with additional optimization from sequence data was able to handle the previously encountered issue of local maximum of the approximated movement function.

## III. Results

Our predictive simulations are shown in Fig. 4. To validate these results, we used the open-source biomechanics dataset by [24], which was created to aid the development of biomechanical models of human locomotion and the design and control of robotic prosthetic legs and exoskeletons. The dataset includes, among other variables, the hip, knee, and ankle joint kinematics of ten able-bodied subjects (age: 30 ± 15 years; height: 1.7 ± 0.9 m; weight: 74.6 ± 9.7 kg). We validated our simulations against subjects walking on level-ground (0° slope) at 1 m/s. Using data from optical motion capture (Vicon, 100 Hz), the joint kinematics were calculated and normalized to percent stride (0-100%) to allow for between and within subject averaging. Compared to the experimental data, our learning algorithm as able to predict the hip, knee, and ankle joint kinematics with total root mean square error (RMSE) values of 8.5°, 10.1°, and 5.7°, respectively. These values are averages for left and right legs. Overall, we were able to simulate biomimetic walking with a 79% degree of similarity compared to the experimental data (Fig. 5).

**Fig. 4.**
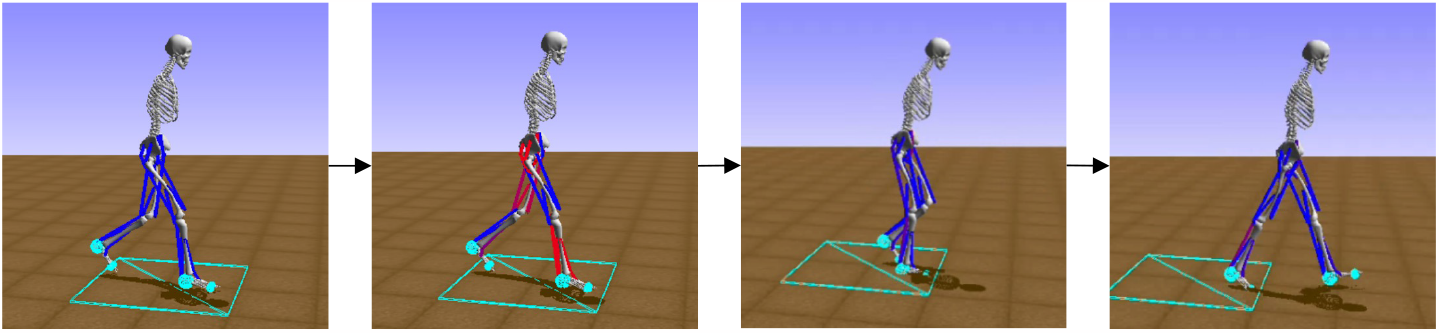
Predictive simulation of human locomotion using our model-free, two-layer reinforcement learning algorithm.

**Fig. 5.**
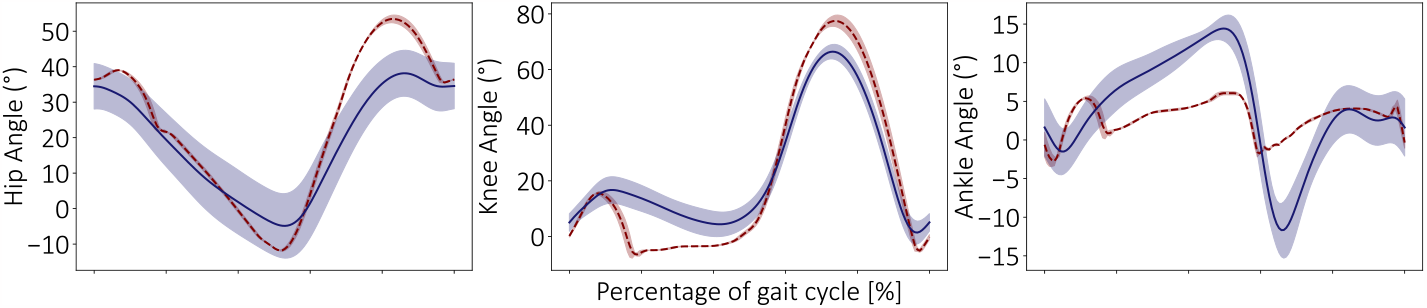
Comparison of hip, knee, and ankle joint kinematics between our predictive simulations (red) and experimental measurements from human walking on level-ground at 1 m/s [24] (blue). The results are averages ± one standard deviation across subjects.

In addition to assessing the predictive capabilities of our algorithm, we also assessed the computational and memory storage requirements in order to support real-time embedded computing for robotic leg control. We compared our requirements with the previous state-of-the-art, a deep reinforcement learning model that won the NeurIPS 2019 Learn to Move challenge [23], which used deep deterministic policy gradient to simulate human locomotion. On average, we required 7 times less RAM and 87 times less CPU compute such that their model required 1,051 MB RAM and an average CPU clock speed of 2,791 ± 14 MHz, whereas our algorithm required only 163 MB and 32 ± 14 MHz, respectively. These comparisons were performed on a machine with an Intel Core i5-8400 CPU @ 2.80GHz and 32 GB RAM.

## IV. Discussion

The goal of this study was to develop a learning algorithm to simulate biomimetic walking that is also suitable for real-time embedded computing for robotic leg control. We proposed a new two-layer Q-learning algorithm, with k-d trees, that supports continuous time series data and a reward model that estimates the degree of muscle activation similarity between the agent and human state-to-action pairs and state-to-action sequences. We used a human musculoskeletal model acting in a physics-based simulation environment [27], [28] to train and evaluate our algorithm to simulate human locomotion. We used imitation learning and artificial biomechanics data to accelerate training via expert demonstrations and used experimental data [24] to validate our simulations. We were able to simulate human walking biomechanics with 79% accuracy compared to experimental data. Moreover, when compared to the previous state-of-the-art [23] that used deep deterministic policy gradient, our algorithm was significantly more efficient with lower computational and memory storage requirements, requiring 7 times less RAM and 87 times less CPU compute. Overall, our method of combining Q-learning with sequential data optimization to train a learning agent for continuous imitation of human locomotion serves as a first step towards the development of bioinspired controllers for robotic prosthetic legs and exoskeletons.

Compared to previous efforts using deep reinforcement learning, our algorithm has several advantages. While deep reinforcement learning has shown potential to control humans and robots in simulation [13], [15], [17], [22], [23], [25], [26], it faces challenges such as limitations in interpretability and large computational requirements. Deep reinforcement learn-ing relies on deep neural networks, which are considered black-box as the internal workings and decision-making processes are not easily explainable [32]. The complex and non-linear transformations applied to the input make it difficult to understand how specific inputs contribute to the output. These models learn representations from data without relying on defined rules or logical reasoning. The decision-making is driven by the learned network weights and activations, which are not readily interpretable as logical rules or symbolic representations. In contrast, our algorithm can be easily interpreted, controlled, and optimized since all data is stored in k-d trees and sequences within the memory buffers. The model includes actions, states, and dictionaries linking actions to states and rewards. It also contains information related to state-to-action pairs and state-to-action sequences. This information enhances our interpretability and model safety, which can be especially important for robotic legs for assistance and rehabilitation of persons with mobility impairments.

Deep reinforcement learning also typically requires large amounts of memory and has computationally intensive operations such as matrix multiplication and convolutions involved in training deep neural networks [32]. These operations require significant processing power and memory bandwidth and are thus often run on high-performance graphics processing units (GPUs) or distributed computing systems with stable electricity. In contrast, our method of updating Q-values via k-d trees is computationally efficient. Q-learning in and of itself is also generally more efficient than deep reinforcement learning as it does not explicitly require a model of the environment dynamics since it learns directly from interactions with the environment. The lack of a model reduces the computational complexity and memory requirements.

However, the development of our algorithm is still preliminary. There were limited vectors in the memory buffer that our learning agent could use during exploitation. The large amount of unused data led to longer searches for the optimal state-action pairs, which increased the computational costs. We also had a limited variety of state-action pairs. This is especially observed in cases where the agent must react to unusual situations like loss of balance or shorter steps. Because it is difficult to simulate these cases, it is hard for our agent to learn them via imitation learning. In addition, our agent lacks adaptability to different movements. Our agent was trained and tested in a stationary simulation environment with a flat surface and no obstacles. This significantly differs from real-world walking environments, which include continuously varying terrains with obstacles and random events [33].

Moving forward, there are a number of opportunities to improve our research. Our algorithm could include a third layer capable of handling transition state-action pairs between different locomotion modes and adding additional instances to our second layer for each mode such that each instance of the second layer is responsible for a specific locomotor task, while the third layer is responsible for task transitions. This hierarchical architecture resembles the control system design of most robotic leg prostheses and exoskeletons [1]. For example, deep learning models of vision [34-35], inertial, and/or EMG data used for high-level locomotion mode recognition could be integrated with reinforcement learning algorithms at the mid-level for end-to-end AI-powered robotic leg control. Also, to support control system design through simulation, another interesting area for future research would be to use large-scale image datasets of real-world walking environments such as [33] to design new photorealistic, physics-based simulation environments, which we plan on exploring.

## Acknowledgment

We want to thank members of the Bionics Lab, part of the Artificial Intelligence and Robotics in Rehabilitation Team at the KITE Research Institute, Toronto Rehabilitation Institute, for their support. This research is dedicated to the people of Ukraine in response to the 2022 Russian invasion.

